# Quantifying shared and unique gene content across 17 microbial ecosystems

**DOI:** 10.1101/2022.07.19.500741

**Authors:** Samuel Zimmerman, Braden T Tierney, Chirag J Patel, Aleksandar D Kostic

**Affiliations:** Department of Biomedical Informatics, Harvard Medical School, Boston, MA 02115, USA; Section on Pathophysiology and Molecular Pharmacology, Joslin Diabetes Center, Boston, MA 02215, USA; Section on Islet Cell and Regenerative Biology, Joslin Diabetes Center, Boston, MA 02215, USA; Department of Microbiology, Harvard Medical School, Boston, MA 02115, USA

## Abstract

Measuring microbial diversity is traditionally based on microbe taxonomy. Here, in contrast, we aimed to quantify heterogeneity in microbial gene content across 14,183 metagenomic samples spanning 17 ecologies including -- 6 human-associated, 7 non-human-host-associated, and 4 in other non-human host environments. In total, we identified 117,629,181 non-redundant genes. The vast majority of genes (66%) occurred in only one sample (i.e. “singletons”). By contrast, we found 1,864 sequences present in every metagenome, but not necessarily every bacterial genome. Additionally, we report datasets of other ecology-associated genes (e.g. abundant in only gut ecosystems) and simultaneously demonstrated that prior microbiome gene catalogs are both incomplete and inaccurately cluster microbial genetic life (e.g. at gene-sequence identifies that are too restrictive). We provide our results and the sets of environmentally-differentiating genes described above at http://www.microbial-genes.bio.

**Importance:** The amount of shared genetic elements has not been quantified between the human microbiome and other host and non-host associated microbiomes. Here we made a gene catalog of 17 different microbial ecosystems and compared them. We show that most species shared between environment and human gut microbiomes are pathogens and that prior gene catalogs described as “near-complete” are far from it. Additionally, over two-thirds of all genes only appear in a single sample and only 1,864 genes (0.001%) are found in all types of metagenomes. These results highlight the large diversity between metagenomes and reveal a new, rare class of genes, those found in every type of metagenome, but not every microbial genome.

## Introduction

The human microbiome, with its established role in modulating host health (1–4), are diverse and arguably a multitude of different ecosystems. Moreover, the microorganisms that inhabit these niches are in no way limited to a single locale across space and time. Many microbes are transient (5, 6) -- passing through the gut, for example, and others, even if colonizing long-term, evolved from ancestors that lived in other hosts or environments altogether (7). The taxonomic, functional, and genetic content of the human microbiome must be contextualized across ecologies to estimate evolutionary history relative to the universe of microbiological life. Public databases that include more than one body site or host are critical in understanding the ecology of microbiomes writ large.

These resources enable answers to basic questions about microbes and their role in human health. Comparisons between human microbiome content -- most frequently the human gut -- and environmental metagenomes have yielded important findings for microbiome ecology in the past, both in terms of the overall taxonomic similarity of ecosystems (at the level of the *16S* gene) as well as the development of its role in host health (8). For example, to this latter point, one group used a combination of microbiome data sources to track the evolutionary history of *Enterococcus* across time, identifying how it adapted to become virulent (9). Other teams have purported the “hygiene hypothesis,” which argues that exposure to certain microbes at a young age can be protective against autoimmune diseases (10). However, testing the hygiene hypothesis requires a thorough understanding of the interactions between environmental and human microbial ecologies.

There is a need to understand the diversity and distribution of bacterial proteins -- as opposed to taxonomies -- across metagenomic life. In this report, we refer to ORFs grouped into clusters by sequence identity and represented by a consensus sequence as “genes”. Some public databases (e.g. Uniprot) (11, 12) contain genes measured across ecologies and environments; however these databases are not designed to address higher resolution comparative biological questions, including enrichment or conservation of specific sequences or functions across different environments. Many other published gene databases focus on single niches -- predominantly gut ecosystems (13).

Additional similar resources tend to focus on taxonomies or genome reconstruction, such as those comprising Metagenome Assembled Genomes (MAGs) (14–16). Compared to MAGs, databases of pan-ecological microbial gene content are lacking. Publications focusing on MAGs have provided insight into the genomic structure of gene content of metagenomes across environments, but by virtue of building a MAG one loses potentially interesting genetic information in genes that are not placeable within a genome. For example, MAGs have been hypothesized to exclude certain bacterial genes (17). While there are recent impressive databases that compare across environments, few, if any, have attempted to explicitly report the “core” genes enriched in different ecosystems (or found in all of them) (16).

Genes, after all, 1) underpin the functional processes of a microbe, 2) encode specific protein products (e.g. antibiotics and other small molecules) of biotechnological importance (18, 19), and 3) have been shown, in our prior work, to outperform taxonomy-based metrics for predicting host phenotypes (4, 20). Finally, a gene-centric view of microbial ecosystems provides one agnostic to the much-debated “microbial species concept” and sidesteps the difficult challenge of quantifying genome “completeness” (21).

Contextualizing the gene universe of metagenomics across ecologies allows one to look into the “conserved” structure of microbial life in terms of genes that are shared between some or all metagenomic ecosystems. Prior work has revealed sets of conserved genes across bacteria (22) and all three domains of life with varying levels of success (23, 24). However, these studies search for conserved genes in a relatively small set of evolutionarily distant organisms. To our knowledge, a search for universally conserved proteins across thousands of metagenomes in different ecologies has not been executed. These hypothetical genes, however, could serve an important purpose through, for example, the “black queen” hypothesis, where functions essential to a community are found not in every cell, but an isolated number of organisms. (25)

Here, we explore the gene universe of metagenomic data, with access to metadata (e.g. host disease state), varying analytic strategies (e.g. genes clustered at different percent identities), and datasets of ecology-specific genes. Specifically, we aimed to quantify the global gene-level (as opposed to taxonomic) similarity between human, other host-associated, and environmental metagenomes, with a focus on 1) measuring singleton content across environments, 2) characterizing and building databases of genes enriched in different metagenome types, and 3) estimating the number of microbial genes that are conserved across every metagenome, but not every microbial genome. We additionally ran our analysis at varying clustering parameters in order to identify the “optimal” clustering identity for gene catalogs.

## Results

### A pan-ecological, gene-centric database of metagenomic life

We predicted microbial genes across 17 different ecologies from 14,183 samples (Table S1). We used publicly available metagenomic samples and assemblies, identifying 1,676,565,730 raw ORFs from assembled contigs. We built a series of non-redundant gene catalogs at varying amino acid identities (from 100% to 30%). At the 30% amino acid identity (see *Methods*), we identified 117,629,181 non-redundant genes (Fig. 1A), which had a total of 15,712 unique protein annotations, 2,628 unique COG annotations, and 2,820 unique EC numbers (Table S2). We quantified the number of non-redundant genes found in each ecology, noting that larger sample sizes, as expected, yielded an increase in the number of genes identified. At 50% identity and 90% identity, we identified a total of 147,534,185 and 254,624,293 non-redundant genes, respectively. All non-redundant gene catalogs are hosted at http://www.microbial-genes.bio. Further, in an effort to provide “environmentally contextualized” data for different genes and metagenomes, we have made available specific gene sets of interest highlighted by the analysis that follows along with host sample characteristics (e.g. host subject age, sex, and disease status). We annotate each gene ID, for example, being conserved across all ecologies or unique to specific ecological subsets.

**Figure 1:**
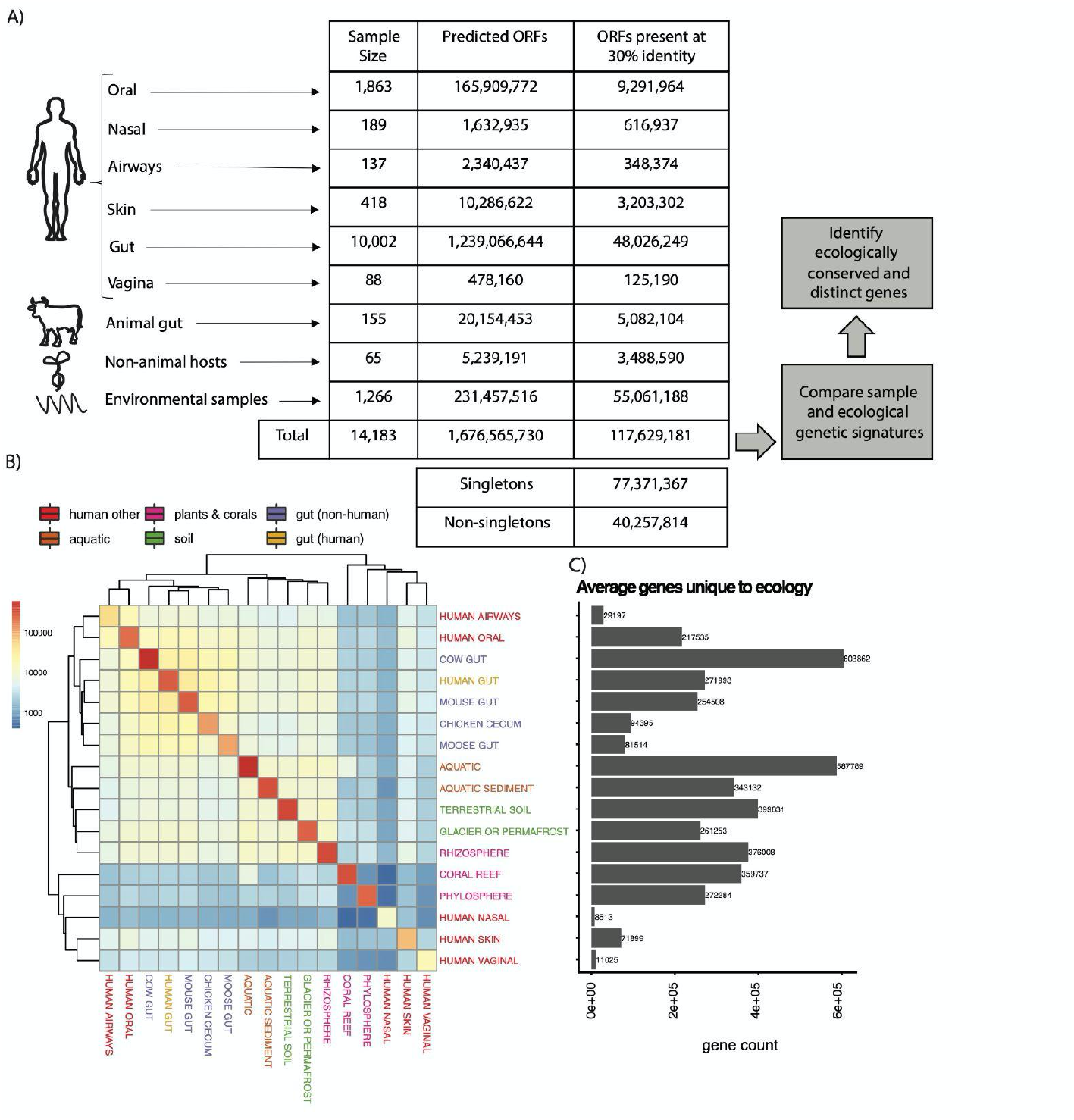
Overview of approach/results and genetic similarity between ecologies. A) Statistics regarding the gene and sample content of our database at the 30% clustered sequence identity and the high-level analytical steps we took in the manuscript. B) Hierarchical clustering on the gene overlap between ecologies as a function of iterative sampling. Each cell represents the average number of genes shared between two ecologies (on rows and columns) after 50 random samplings. Cell color is in units of number of genes. Color of text corresponds to broader ecology class. C) The average number of genes unique to (found only in) a given ecology. This number corresponds to the values in the diagonal.

We counted the proportion of “singleton” genes (26). Even with a lower percent identity cutoff (30% compared to 50%) and a more conservative clustering algorithm, we found that the proportion of singleton to non-singleton genes increased (66% vs 50%), with 77,371,367 singletons and 40,257,814 non-singletons compared to our previous database (26). This indicates that at lower percent identities over half of the resulting gene clusters are still singletons, despite a greater number of samples and ecosystem types than in prior analyses.

### Using microbiome genetic fingerprints to estimate the size of as well as similarities between and within microbial pangenomes

Akin to prior efforts working at the taxonomic level, we aimed to compare the similarity of different ecologies based on gene content (i.e. the presence or absence of a gene assembled in a given sample) in an effort to determine, overall, how pan-genomically similar different environments are. To control for bias due to sample size and the sequencing depth of individual samples, we iteratively compared the overlapping genes between pairs of samples from different ecologies (see Methods), estimating the overall average number of genes unique to and shared between samples. Ecologies segregated, broadly speaking, according to broader environmental context, including gut-associated samples, environmental samples, and non-human hosts (Fig. 1B). For example, gut microbiome samples tended to group together (i.e. from chicken, moose, mouse, human, and cow). Analogously, environmentally-sampled, non-host-associated microbiomes (e.g. aquatic, aquatic sediment, terrestrial soil) had similar overlapping gene content. Samples from coral reefs, the phyllosphere, the human nasal microbiome, and the human vaginal microbiome shared limited gene content with other ecologies. Nasal ecologies, on average, share the fewest genes with any other ecology (Fig. 1B). Samples from human gut and mouse microbiomes shared the most genes with other ecologies on average across all sampling iterations.

We found that the number of unique genes in an ecology did not correlate to the sample size for that ecology. Further, to confirm that our results were not an artifact of our method, we performed a permutation test, randomizing sample gene content and ecology, showing that there was limited similarity between the values in the heatmap in Fig. 1B and this randomized dataset (Fig. S1).

We additionally counted the number of genes that were unique to a given ecology compared to all others (Fig. 1C). A number of environments contained limited numbers of unique genes. These included the human nasal cavity (average unique gene count = 8,613, sample size = 189), human vaginal samples (average unique gene count = 11,025, sample size = 88), human airways (average unique gene count = 29,197, sample size = 137), human skin (average unique gene count = 71,899, sample size = 418), chicken cecum (average unique gene count = 94,395, sample size = 27), and moose gut (average unique gene count = 81,514, sample size = 4). Conversely, the cow gut microbiome had the highest number of genes on average unique to any given ecology (average gene count = 603,862, sample size = 24). In other words, the cow microbiome is genetically more “distinct” from all other microbiomes, on average, than the vaginal microbiome, for example.

The analyses in Figure 1B-C, considered together, revealed insights into the comparative genetic content of different ecologies. For example, the vaginal microbiome had little overlap with other ecologies in terms of genetic content (Fig. 1B). However, any pair of vaginal samples were also highly dissimilar genetically. In other words, two samples from the vaginal microbiome will have little in common with other ecologies as well as each other. By comparison, at the taxonomic level, the vaginal microbiome has ostensibly low phylogenetic diversity (27). Therefore, this analysis indicates that the vaginal microbiome has low alpha diversity yet high beta diversity. We found similar results for the nasal microbiome and the skin microbiome. However, the opposite is true for other ecologies, such as the chicken cecum and mouse gut samples, which had high numbers of shared genes and lower “uniqueness,” potentially indicating increased consistency in species pangenomes.

We additionally carried out unsupervised clustering analysis, identifying and taxonomically characterizing 7 discrete gene clusters (Text S1). We reproduced the result (previously shown at the taxonomic level) that metagenomes segregate according to ecosystem, while also demonstrating that health status does not shape metagenomic gene content compared to westernization status and age (Fig. S7,S8).

### Ecologically contextualizing the gut ecosystem yields a database of discrete and shared gene sets

We next aimed to contextualize the gene content of the human microbiome by identifying ecology-specific (e.g. broadly gut associated) genetic elements of metagenomics. We identified sets of genes that, compared to all other ecologies, were abundant in 1) human gut and environmental samples 2) human gut and non-human gut samples, and 3) environmental samples alone. Specifically, we identified genes that were prevalent in these categories in our initial gene catalog and then tested their differential abundances between different environments in an independent set of 422 samples from our ecologies of interest (see *Methods,* Table S3). We identified 59,944 enriched genes in the human gut and environmental samples, 117,443 genes enriched in the human gut and non-human gut samples, and 39,623 genes enriched in the environmental samples (Fig. 3A).

We found functionally and taxonomically distinct sets of genes in each of the three comparisons we tested (Fig. 2, left column). We additionally explored the specific ecologies in which differential genes overlapped (Fig. 2, right column). For example, the genes overlapping between gut samples and environmental samples were generally found in ecologies containing sediment (e.g. terrestrial soil, aquatic sediment), and non-human gut samples. The human gut stood alone (Fig. 2B), generally speaking. Finally, we found that gene abundance in the environmental samples tended to be shared between samples containing sediment, as opposed to purely aquatic samples (Fig. 2C).

**Figure 2:**
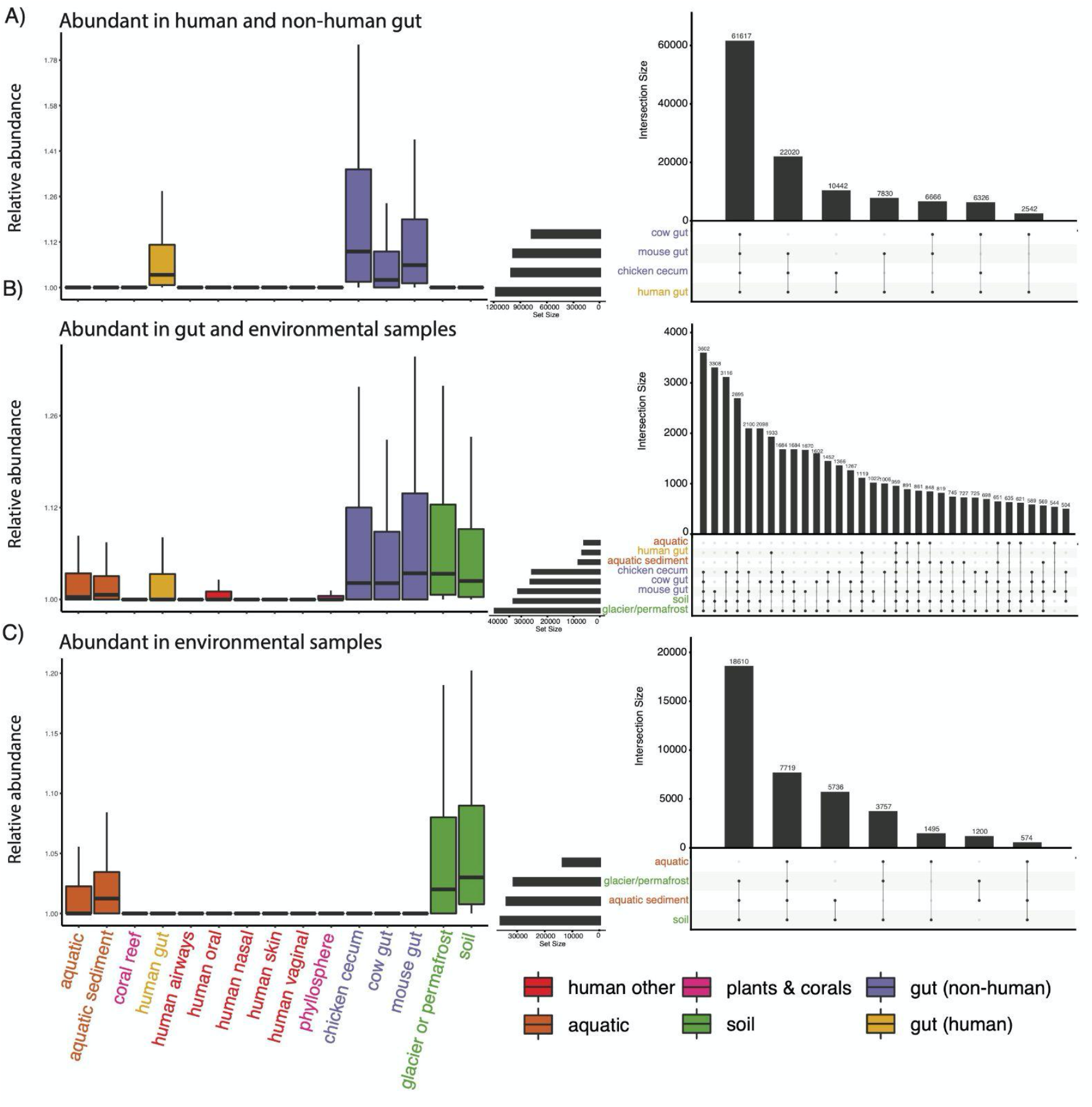
Genes abundant in intersecting ecological groups. Left column: The abundance of genes found to be significantly differentially abundant (in a separate cohort) in the described sample types. Right column: The number of specific overlaps in abundant genes between specific ecologies.

We identified the COG categories of each annotated gene, quantifying the proportion of genes in each category (Fig. 3B). Carbohydrate processing was the top category for both human gut/non human gut genes (∼20% of all genes with annotations) and human gut/environment genes (∼15% of all genes with annotations). Genes abundant in both environmental samples and environmental/gut samples both contained many annotations to the broad categories of energy production and conversion and signal transduction mechanisms.

**Figure 3:**
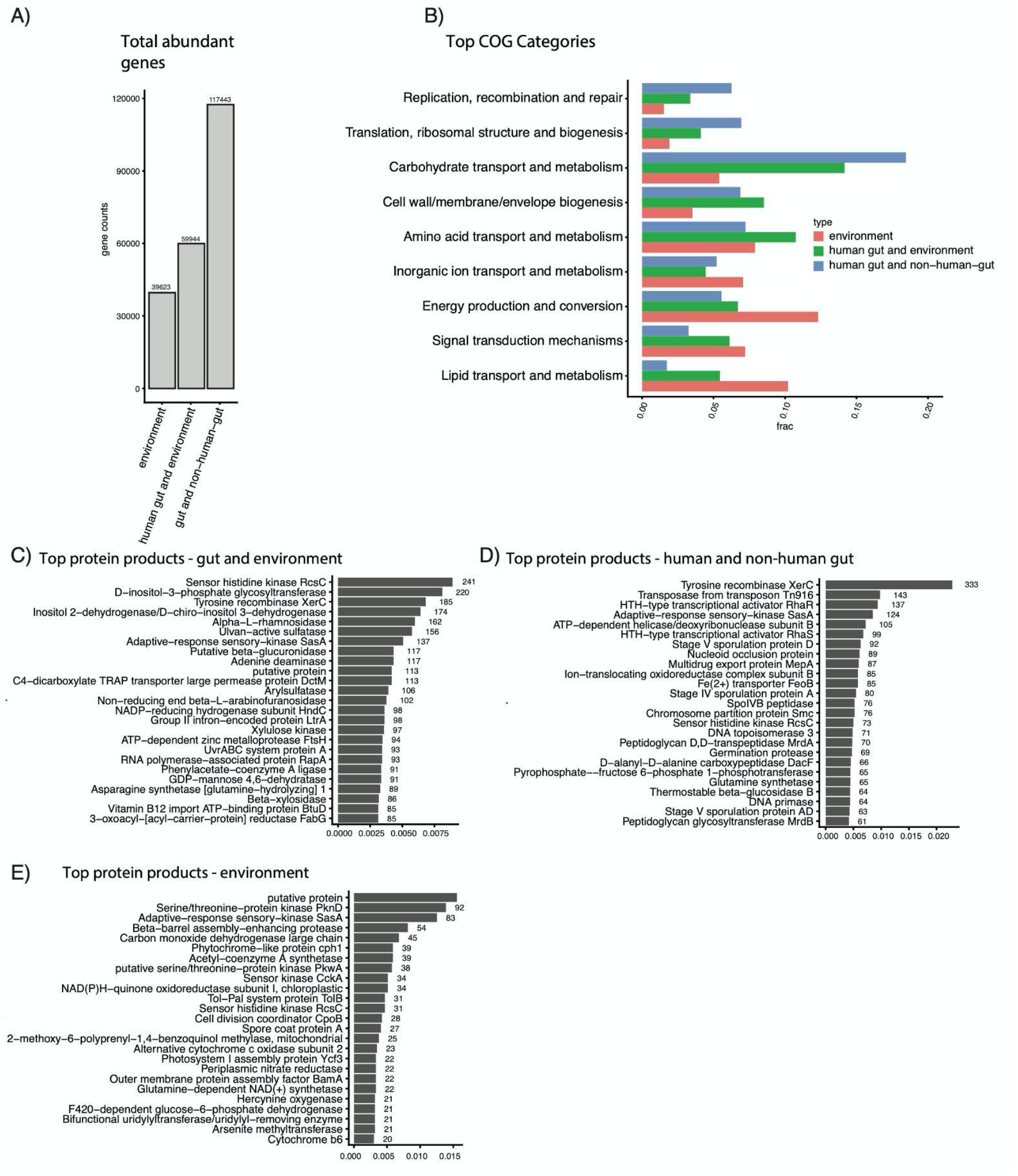
Functional analysis of genes abundant in different ecological contexts. A) The total number of genes captured in the differential abundance analysis. B) The fraction of genes in the different high-level COG categories, by intersection. C-E) The top 25 most common protein products for each ecological comparison considered.

We additionally analyzed the most common protein annotations in our differentially abundant genesets, finding that each set contains functionally distinct proteins (Fig. 3C-E). These functions did not originate from specific bacterial taxonomies, but rather were represented across the phylogenetic spectrum (Fig. S9). Many of the most common proteins in the environmental category (Fig. 3E) were involved in photosynthesis (e.g. photosystem, cytochrome, and phytochrome proteins) and environmental nutrients (e.g. nitrate reductase, arsenite methyltransferase). Genes abundant in both the gut and environmental ecologies (Fig. 3C) were associated with a variety of nutrient import and export functions (e.g. Vitamin B12 import ATP−binding protein BtuD, C4−dicarboxylate TRAP transporter large permease protein DctM). The top two proteins abundant in human and non-human gut ecosystems (Fig. 3D) were both associated with horizontal gene transfer (tyrosine recombinase XerC, transposase from transposon Tn916). Other annotations in this category were associated with sporulation (e.g. stage V sporulation protein AD), antibiotic resistance (e.g. multidrug export protein MepA), and cell wall construction (e.g. peptidoglycan glycosyltransferase MrdB, Peptidoglycan D,D−transpeptidase MrdA). We were unable to find genes that were mutually and significantly abundant in all 6 human body sites queried compared to other ecologies (Fig. S10).

### Ecologically contextualized metagenomic architectures identify the genetic features underlying the multi-ecosystem capability of pathogens

As an application of the ecologically differential genes we identified, we next sought to identify the taxonomic annotation of genes that spanned environments. We hypothesized that genes found in specific or multiple environments would be those potentially required for an organism to survive in said ecosystems, warranting their further investigation in *in vivo* studies. We compared the most common taxonomic annotations of genes that were abundant in both the gut and at least one of the other ecological categories. First, we taxonomically annotated genes that were prevalent and abundant in the same three comparisons as in Figure 2. Despite the difficulty of annotating genes to taxonomy (e.g. some annotations, like “*clostridia bacterium*” being broad and not informative on their own), visualizing the resulting taxonomic bins displayed a clear pattern in phylogeny across human, non-human, and environmental ecologies (Fig. 4). Human gut and non-human gut samples overlapped predominantly in the *Firmicutes* phyla. Very few *Firmicutes* annotations were highly abundant in environmental samples, relatively speaking. The exceptions to this were members of the family *Christensenellaceae*, which had genes annotated as abundant in all ecologies. This was also the case for *Roseburia faecis*, which was the most prominent member of the *Roseburia* genus that had many genes in both gut and environmental ecologies. On the other hand, organisms of the phyla *Bacteroides* tended to straddle gut-associated ecosystems and terrestrial ecosystems, but had limited ostensible genetic representation in aquatic ecosystems. *Proteobacteria* tended to characterize non-human-gut and environmental samples, with certain organisms, like *Vibrio cholera*, having representative genes present across all ecologies. We noted that pathogenic microbes, like *V. cholera* or *Mycobacterium tuberculosis*, stood out in the sense that they spanned ecologies.

**Figure 4:**
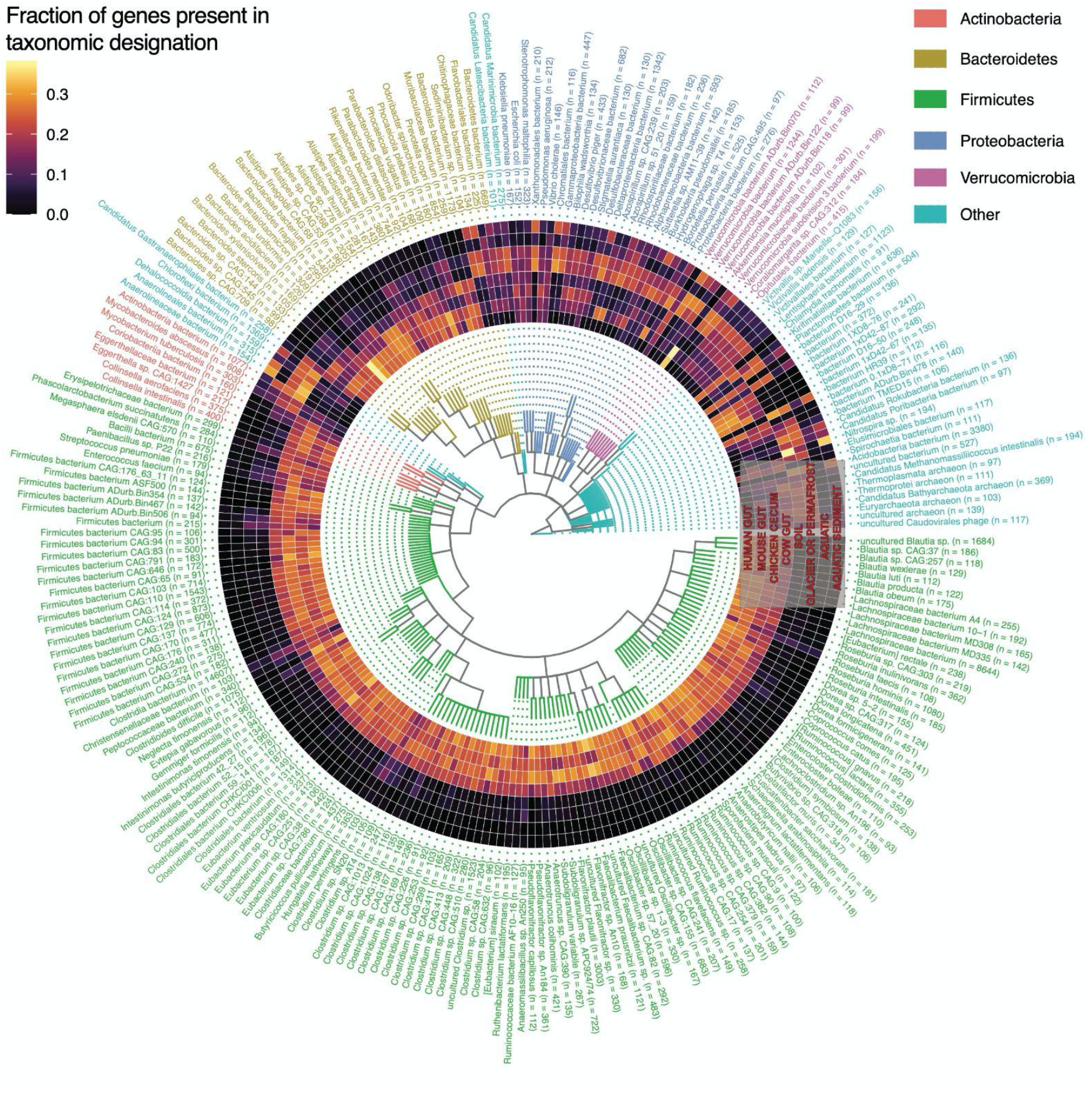
Taxonomically contextualizing the genetic content of the human gut microbiome. Each ring corresponds to a different ecology. Each “row” corresponds to a different taxonomic annotation for a consensus gene that was abundant in the human gut microbiome and at least one other environmental or non-human-gut system (as observed in Figure 3-4). The colors correspond to the fraction of all genes with a given annotation that were indicated as abundant in a given ecology. Text color corresponds to phyla.

The majority of overlapping taxonomic features across host-associated environments and free-living ecologies occurred in non-human gut samples and samples containing sediment. We did not identify any taxonomic annotations that were only found in the human gut, however, we did notice that certain species, like *Bacteroides xylanisolvens* and the genus *Faecalibacterium* were notably prevalent in a subset of gut samples (human and mouse/human and chicken, respectively).

Next, we examined the taxonomic distribution of genes that were differentially abundant in the 6 human-associated ecologies in our dataset compared to environmental, plant, and coral ecologies (Fig. 5). The result demonstrated the limited genetic overlap of human microbiome ecologies, notably the vaginal, skin, airways, and nasal microbiomes with the gut and oral ecologies. However, we did find that the oral microbiome shared the most genetic similarities with all other body sites, most noticeably but not limited to the gut, where it overlapped predominantly in terms of *Firmicutes* and *Bacteroidetes* annotations. The vast majority of species tended to be predominantly site specific, such as *Cutibacterium acnes* in the skin microbiome or many of the *Streptococci* in the oral microbiome. Species that straddled multiple sites (>3) and/or had many (>100) genes annotated to them, tended, like before, to be potentially pathogenic. These included *Staphylococcus aureus, Acinetobacter baumannii, Cronobacter sakazakii, and Enterococcus faecalis*.

**Figure 5:**
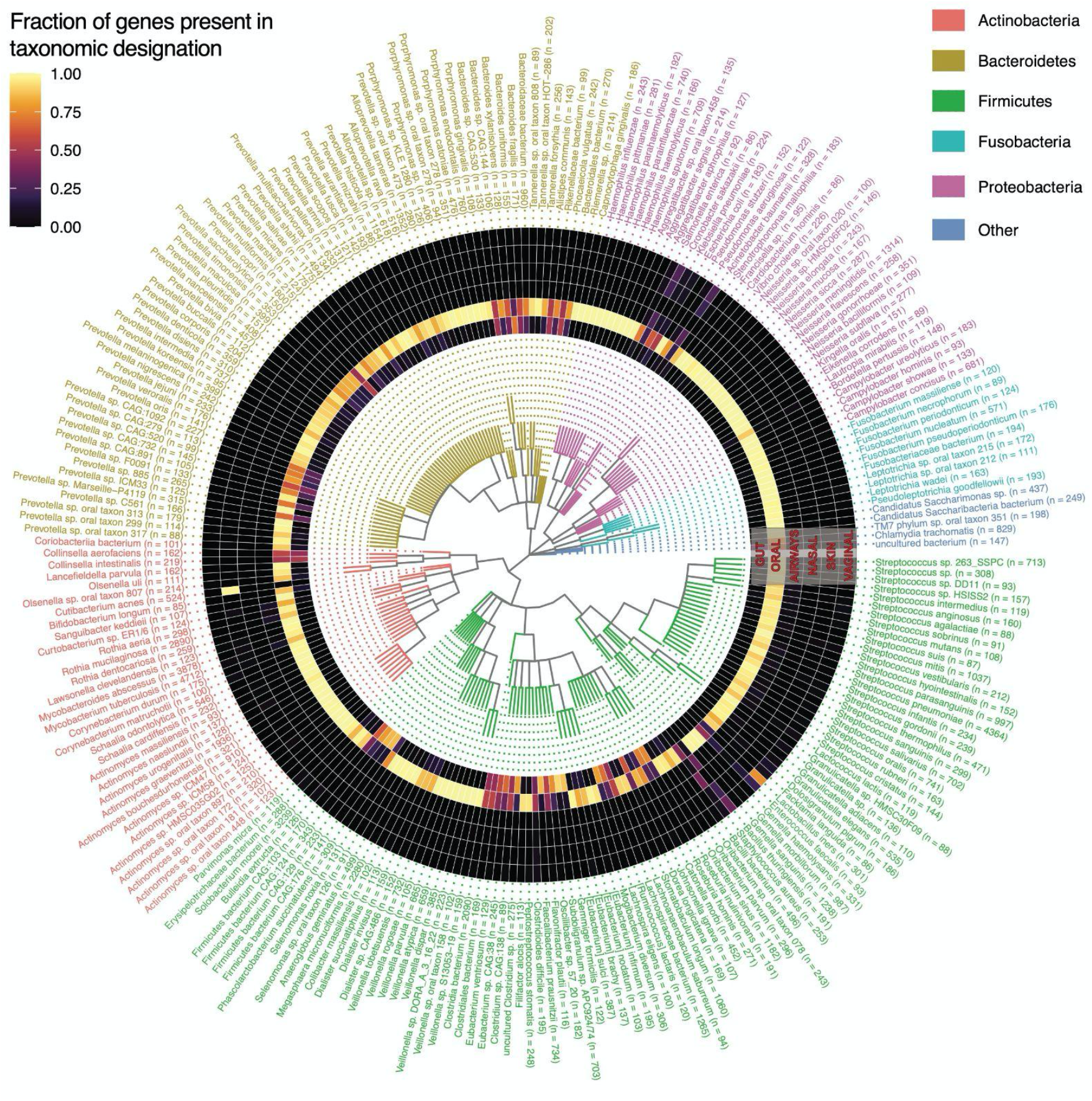
Taxonomically contextualizing the genetic content of the global human microbiome. Each ring corresponds to a different ecology. Each “row” corresponds to a different taxonomic annotation for a consensus gene that was abundant in at least two human microbiome body sites. The colors correspond to the fraction of all genes with a given annotation that were indicated as abundant in a given ecology. Text color corresponds to phyla.

### At least 1,864 genes -- not identifiable at high clustering identities -- comprise the ecologically conserved elements of metagenomics

Finally, we sought to identify sequences that were prevalent and abundant in all 17 ecologies in our database. At 30% clustering identity, we identified 1,864 genes that were found on contigs in at least 1 sample for all ecologies, and we quantified the abundance of the 1,864 genes in the same 422 independent samples from the prior analysis (Fig. 6A, Table S4), finding that both prevalence (median = 0.84) and abundance varied substantially as a function of ecology. We additionally compared their prevalence to that of the bac120 genes used to taxonomically classify bacteria in the Genome-Taxonomy-Database (GTDB), which are derived from specifically single copy genes present in at least 95% of a database of microbial genomes (not metagenomes) (28). We hypothesized that searching for globally conserved genes with our approach would be sufficient to recover these genes. We found this to be the case. The bac120 genes had a slightly higher overall prevalence (median = 0.96), however we were able to identify (via alignment) 114/120 (95%) in our 1,864 genes (Fig. 6B). Notably, the 10 most prevalent genes did not align to the bac120 set.

**Figure 6:**
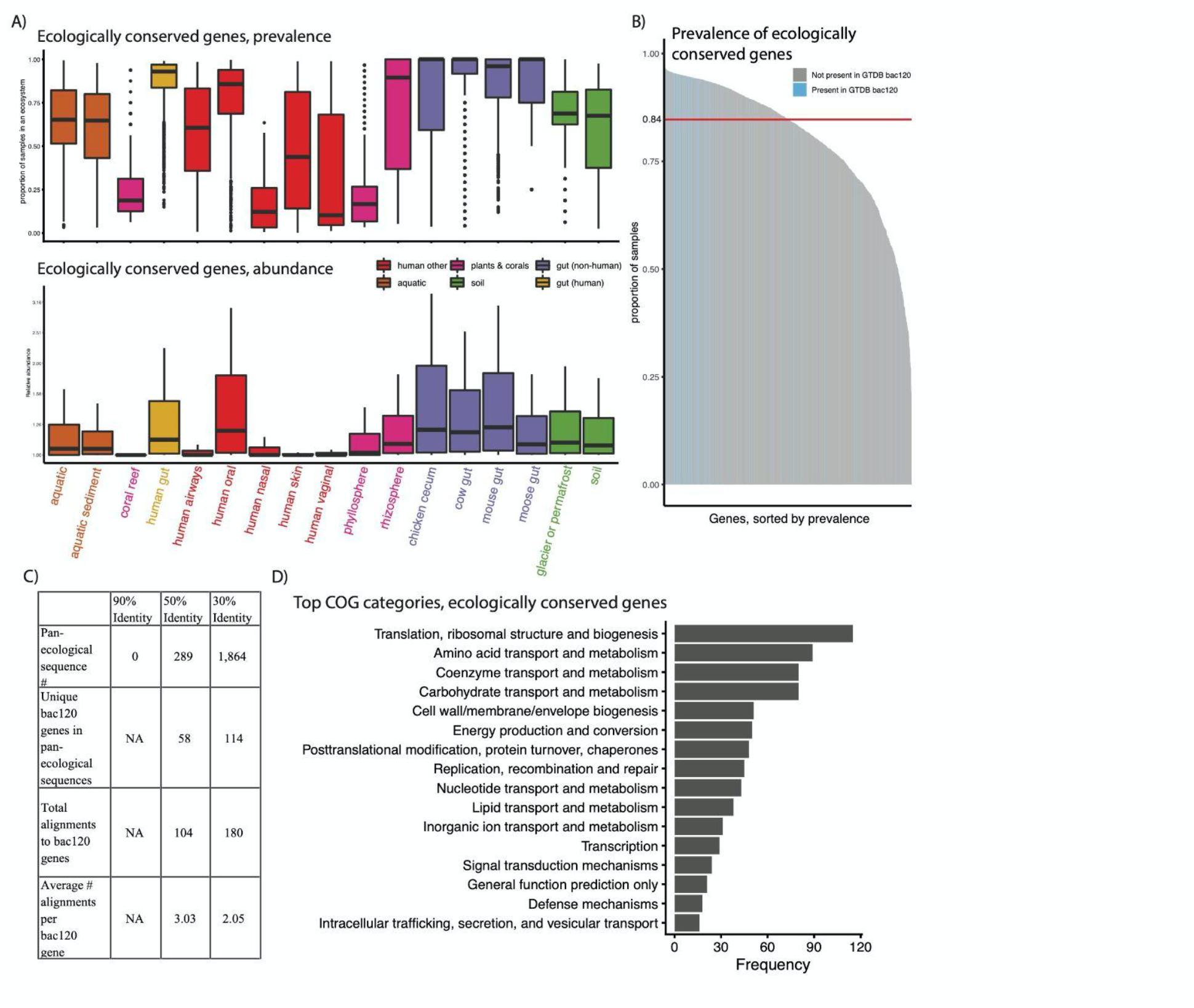
The pan-ecologically conserved genes of metagenomics. A) The prevalence and abundance (in the 422 external samples used in prior figures), by ecology, of all 1,864 genes found to be assembled at least once in all 17 ecologies. B) The prevalence of all 1,864 ecologically conserved genes, with colors corresponding to whether or not they aligned to a GTDB bac120 gene. C) For different percent identities, the number of pan-ecological sequences, the number that align to GTDB bac120 genes, and the average number of bac120 genes aligning to a given identified conserved gene. D) The most common COG category annotations for the 30% identity ecologically conserved genes.

We additionally compared the number of conserved genes found at 30% to those found at the 90% and 50% amino acid clustering identities (Fig. 6C, Fig. S11). We found at 90% identity there were no genes found in every ecology, indicating that 90% is too strict a clustering cutoff, as at least the bac120 genes should be identified in the majority of samples. Similarly, we only identified 289 ecologically conserved genes in our 50% gene catalog, however these aligned uniquely to only 58 of the bac120 genes (48%), and each bac120 gene aligned to an average of 3.05 sequences (compared to 2.03 at 30% identity) indicating that genes clustered at 50% are, on average, “under-clustered” for this particular use-case.

We next examined the function of these “ecologically conserved” genes (Table S4). We hypothesized that perhaps they contained functional redundancy, and that we were in fact referring to genes as “different” when in fact they did the same thing. We found this not to be the case. While we did find that the bac120 genes aligned to 180 of out 1,864, indicating mild functional redundancy, we found that, after removing 600 hypothetical proteins, the remaining 1,264 genes were annotated as coding for 1,165 unique proteins.

While, overall, the 1,864 proteins we identified were conserved across microbial life (Fig. 6A), we sought to identify if, specifically, these genes reflected functions not found in the bac120. We found (Fig. 6D), that while the GTDB genes tended to reflect what one would hypothesize to be highly conserved metabolic processes (e.g. ribosome-associated-genes), the broader set of ecologically conserved genes did not reveal any consistent set of functions, including a variety of processes from nutrient transport to cell-to-cell signaling. We hypothesize that these may comprise “ecologically core” functions, those not found in every microbial cell, but rather in all metagenomes.

## Discussion

The genomes of microorganisms that colonize the human ecosystem contain a long evolutionary history spanning many hosts and environments. As a result, to understand the evolutionary and functional dynamics that drive the human microbiome -- an ecosystem that modulates so many aspects of human health -- we must consider the full ecological spectrum in which microorganisms are found. Our approach here specifically relies on attempting this goal by comparing Open-Reading-Frame presence or absence in order to genetically contextualize and compare metagenomic life.

We incorporate metagenomes from 17 different ecologies, comparing their genetic content, and providing sets of genes that we found to be prevalent and abundant in different ecological contexts, with a specific focus on the gut microbiome. This dataset, with its focus on differential distribution of genes, instead of genomes, complements existing published resources. Further, given that many of its samples are shared between our resource and others, these datasets could feasibly be linked into one, meta-database, where genes can be linked to genomes effectively. For example, our identification of pathogen-associated genes that span environments could be identified in pathogen MAGs and further explored as potential mechanisms of pathogen adaptability to varied environments.

Outside of challenges raised with all gene catalog analyses (e.g. sub-optimal clustering algorithms) (29), one potential drawback of this analysis is the variation in sample sizes, assembly quality across environments (e.g. corals vs human gut samples), and sequencing depth across environments which could bias our results in terms of the unsupervised UMAP clustering and the differential abundance analysis. Future work should aim to increase sampling from all metagenomic ecologies in order to fully explore the gene universe of microbiome writ large. We additionally note that there is no known “biologically correct” percent identity cutoff for gene catalog analysis, and certainly some genes likely diverge more than others. Therefore, there is a need for additional algorithmic development that builds functionally-motivated gene clusters using different or additional metrics to percent identity alone.

On this note, however, another finding of our report is the utility of clustering at 30% amino acid identity, as opposed to the higher levels reported in the literature (30). Clustering based on sequence identity has been contentious in the literature, and clustering at lower percent identities has been computationally infeasible until the development of recent algorithms (29). The inability to identify functionally discrete genes at other clustering identities indicates that prior gene catalog analyses could be improved -- or at least better understood -- but applying slightly different algorithmic approaches. While we do provide 90% and 50% amino acid identity gene catalogs in our online results, we chose to report the majority of the analysis in this paper using our 30% identity dataset. This is because 1) homologous proteins are known to diverge up to 30% identity and 2) at least one other database additionally clusters at 30% and reports high quality clusters with homogeneous functions (11, 31). However, despite the increased stringency of our clustering analysis over past work, we were still able to reproduce a striking prior result: the dominance of singleton genes in metagenomes. When clustering at 30% identity with a more conservative (less greedy) algorithm than before, we still found that singletons, genes assembling in only one sample, comprised >50% of our gene catalog. Given our prior efforts demonstrating that singletons are unlikely to be false positive genes (26), this implies once more that the gene universe of bacteria *writ large* exceeds what we possibly could have imagined.

Clustering at 30% identity enabled us to identify and characterize 1,864 microbial genes that were ecologically pervasive. We hypothesize, specifically, that these genes represent a class of genetic elements with functions present not in every microbial genome, but rather in every metagenome, implying perhaps that they carry out functions potentially fundamental to community structure. Further, the fact that these ∼1800 genes did not overlap substantially in function increased our confidence in our choice of a 30% clustering identity. We also showed that these genes would not be detectable using other clustering identities deployed in the literature, 95% nucleotide-level identity,(11, 13, 30) compared to a more aggressive threshold, 30% amino-acid identity.

Further, the identification of both singletons as well as conserved genes at 30% amino acid clustering identity indicates that prior gene catalog analyses do not even approach capturing the entire genetic diversity of metagenomic systems. We identified 48 million (Fig. 1) genes in the gut microbiome alone -- many more than previously observed, and we do expect this number to continue to increase significantly as more humans are sequenced. This calls into question the claims made by prior work that previous sets of microbial gene catalogs of the human gut microbiomes were “close-to-complete.”(13) Notably these claims were made using 95% nucleotide identity first with a 3.3 million gene catalog (30), then again with a 9.9 million gene catalog (13). It may be that “complete” databases of microbial life -- whether they be at the gene or strain level -are infeasible. That said, each of these studies has provided a different window into metagenomic ecosystems, and their expansion -- as we attempt to do here -- will hopefully continue to do so, whether or not we are able to “completely” sequence the gene content of microbial life.

## Methods

### Overview of approach

We aimed to compare the gene content of metagenomes across different ecological contexts and provide a database where genes specific to and shared across environments can be queried. We built a non-redundant gene catalog of 14,183 samples from 462 studies across 17 different ecological contexts. We compared the gene content of each of these ecologies, identifying which are most similar and different to each other. We then used a dimensionality reduction approach to cluster samples on the genetic level, identifying distinct ecological, taxonomic, functional, and phenotypic features of each cluster. Based on prevalence, we identified the “ecologically core” elements of metagenomes. We compared these to a database of known conserved genes in bacteria (bac120 from the Genome Taxonomy Database). We additionally used differential abundance analysis on 422 separate samples to identify genes that were abundant in different ecological sets (e.g. in gut-associated-biomes). We provide a database of our results at http://www.microbial-genes.bio.

### Data aggregation

We aggregated publicly available data from a variety of sources, including already-published, assembled metagenomes as well as raw shotgun sequencing read data. A full list of data used and summary statistics (e.g. number of genes per sample) is present in Table S1. In total, after filtering for samples with low numbers of genes (see next section), we aggregated 14,183 samples from 462 different datasets/studies.

For the human-associated samples, we aggregated assembled samples from our prior work (26) and also one published database that contained assembled contigs (but not predicted genes) (15). This ended up yielding 12,697 samples from 6 different human body sites and 65 studies.

For the environmental and non-human-host samples, we aggregated data from the MGnify database (https://www.ebi.ac.uk/metagenomics/). We selected all non-human host-associated and environmental, assembled metagenomes and accessed their download links. We then manually curated the ∼2200 samples (accessed in winter 2020), removing 1) duplicated assemblies and 2) samples where we could not clearly confirm the source of the metagenome by identifying the publication or raw upload data. We found a substantial number of duplicated and/or mis-annotated samples, so our final sample tally came to 1,486 samples from 397 datasets from 11 ecologies.

### Gene catalog construction

For the 1,694 samples that we assembled ourselves (i.e. were not already assembled by others), we removed adapter contamination using bbduk, from bbtools V37.62, removed contamination from the human genome with bmtagger V3.101, and carried out *de novo* assembly with MetaSPAdes (32) V3.13.1 using the default parameters. When MetaSPAdes failed due to a currently unresolved bug, we used MEGAHIT V1.2.9 (parameters: --mem-flag 2,) (33). We measured N50s with assemblies using MetaQuast V5.0.2 (parameters: --max-ref-number 0 -t 15 --fast -m 0). Note that there were 6 samples that, after we downloaded them and included them in our database, became inaccessible (e.g.https://www.ebi.ac.uk/ena/browser/view/ERZ781377), and as a result we were unable to compute N50s for them. However, all of these samples had high gene counts (>1,000) and originated from ecologies for which we had many samples (aquatic, cow, mouse) and as a result we are not concerned about assembly quality nor their effect on our results.

To predict Open Reading Frames (ORFs) from assembled contigs, we used Prokka (34) V1.14.6 (parameters: --force, --addgenes, --metagenome, --mincontiglen 1). The functional annotations (e.g. protein names, COGs) in this manuscript are those automatically generated by Prokka. We removed samples that contained fewer than 100 genes. In our prior database we removed genes <100bp long, however given recent work by other teams indicating the physiological import of such short sequences, we chose to leave them in our study (35).

We then clustered all Open Reading Frames predicted by Prokka using the linclust (36) and cluster functions from the MMseqs2 (37) package. As in previous work (26), we initially iteratively clustered (using linclust) our Open-Reading-Frames, starting at 100% amino acid identity and decreasing, in increments of 5%, down to 30% identity (other parameters: -c [alignment coverage] 0.9). In order to further increase the clustering of the gene catalog we used for the majority of analysis in this paper, we used the more conservative “cluster” function (same parameters as before at 30% identity) in order to reduce the total number of genes in our 30%, 50%, and 90% catalogs. We chose to use the 30% gene catalog for our analysis due to 1) the use of a 30% cutoff in the literature and 2) the fact that many genes -- even those involved in core processes -- contain homology at near 30% level (11, 31). We provide these 50%, 90%, and 30% gene catalogs on our resource.

### Gene-level taxonomic annotation

We assigned best-possible taxonomic annotations to each gene of interest using the built-in Least Common Ancestor computation tool of the Diamond (38) aligner with the default parameters. In doing so, we aligned our 30% gene catalog back to the NR reference database and NCBI taxonomy data, annotating, where possible, the NCBI taxonomy of each gene.

### Comparative subsampling of metagenomic ecologies

In Figure 1B-C, we used a subsampling strategy to estimate the overall genetic similarity between different ecologies. This was to control for variation in sample size and sequencing depth, as the ecologies we sampled had radically different numbers of observations (e.g. a minimum of 4 moose samples to a maximum of 10,002 human gut microbiome samples). Over 50 iterations, we took 1 sample from each ecology. We counted the number of ecologies each 30% consensus gene appeared in (a minimum of 1 and a maximum of 17). We repeated this process 50 times and averaged the number of shared genes between each pairwise set of ecologies, as well as the average number of genes that were unique to each ecology. We additionally took this approach for our permutation test (Fig. S1), which we did as a point of comparison, randomizing the gene-sample-ecology mapping.

### Gene-level dimensionality reduction and clustering of samples

We aimed to cluster samples based on gene presence-absence using a series of dimensionality reduction approaches. This is a computationally challenging problem due to the massive dimensionality: a sparse matrix with 15 million genes and 14,183 samples. To address this we performed Latent Semantic Indexing (LSI), a dimensionality reduction method used in other fields like single cell RNA-SEQ (Cusanovich et al. 2015). In this approach, we first perform term frequency-inverse document frequency (TF-IDF) normalization on a binary “presence absence” matrix. This is a two step process where we first divide the presence or absence of a gene (1 or 0) by the total number of genes in each sample. This is called the term frequency (TF) and normalizes by the number of genes per sample to make each sample comparable with each other. Next, we calculate the inverse document frequency (IDF), which is the log of the number of samples (plus 1) divided by the total occurrences of each gene in all samples. This effectively increases the weight of rarer genes that may distinguish a sample from others. Lastly, we use matrix multiplication to multiply TF by IDF. This procedure normalizes across samples to correct for differences in gene number and across genes to give higher values to more rare genes.

In the second step of LSI, we input this matrix into a Singular Value Decomposition (SVD) to reduce the dimensionality of our dataset into 10, 50, or 100 principal components. We then used UMAP (39) to further reduce our output to two plottable vectors. UMAP was run on each dataset 12 times with a combination of different parameters specified below. Reducing the output to two vectors with UMAP facilitates clustering by making the density of samples in a dataset more evident. This allows density based clustering algorithms such as DBSCAN to more easily cluster data. Next, we ran DBSCAN on each UMAP output four times, varying the parameter that specifies the minimum number of samples in a cluster. The clustering results that maximized silhouette score while minimizing the number of unclustered samples were used for downstream analysis and interpretation (Fig. S3).

As described above, we tried a number of different parameters for the SVD, UMAP, and DBSCAN components of our clustering pipeline in order to test the robustness of our clusters as well as select the “best” parameters (SVD: 10, 50, and 100 single values [SV]; UMAP: n_neighbors=10, 15, 20, and 30, and min_dist=0.1, 0.25, and 0.5, and HDBSCAN: minPts=25, 50, 75, and 100). We used the silhouette score and number of unclustered samples to optimize the parameters. We found that the following parameters optimized silhouette score and number of clustered samples (SVD: singular values [SV]=50, UMAP:n_neighbors:10, min_dist=0.1, HDBSCAN: minPts=50, n_components=2), and that the clusters were robust to variation in parameters (Fig. S3). R version 4.0.3, package RSpectra V0.16 was used to perform SVD. R version 4.0.2 was used to do the UMAP and clustering. R packages umap V0.2.7.0 and dbscan V1.1-6 were used to do the clustering.

### Dimensional Reduction of UMAP gut-associated clusters

After clustering all samples we performed additional Latent Semantic Indexing solely on the samples in the clusters containing human gut microbiome metagenomes (clusters 2, 3, 4, and 7). As above, we first used TFIDF followed by SVD on the gene presence-absence matrices. Then we used UMAP to reduce our output into a two dimensional plane. The parameters used for reclustering all subclusters were SVD: SV=50, UMAP:n_neighbors:10, min_dist=0.1, HDBSCAN: minPts=50, n_components=2).

### Cluster-based functional and taxonomic enrichment analysis

We used the taxonomic and functional annotations of the consensus genes in our 30% amino acid identity catalog to identify those that were enriched in samples present in each of the 7 clusters identified by our prior clustering analysis. To achieve this end, we used Chi-Squared tests, populating a contingency table with the number of genes that have the protein-name or taxonomic annotation of interest within a cluster and outside of a cluster as well as the overall number of taxonomic/functional annotations within and outside of a given cluster. Enriched samples were those with a BY adjusted p-value < 0.05. Fisher’s exact tests were used when an element of the contingency table had less than 6 observations.

### Identification of ecologically conserved genes and differentially abundant genes

We identified “ecologically core” genes by finding those that were present at least once in each of our 17 ecologies.

We used both prevalence and abundance to also find genes that were 1) shared between the gut microbiome and other non-human gut microbiomes and 2) shared between the gut microbiome and terrestrial/aquatic microbiomes, and 3) shared between terrestrial and aquatic microbiomes only. We took 1) genes prevalent in at least one sample of the ecologies of interest, 2) but not present in any other samples, and 3) those that had statistically significantly higher abundance than all other samples (BY adjusted p-value < 0.05)

### Comparison of ecologically conserved genes to the Genome-Taxonomy-Database (GTDB) bac120

We gathered the HMMs corresponding to the 120 GTDB bac120 genes in Parks et al. 2017 from Pfam (V 33.1) and TIGRFAM (V 15.0). We used hmmsearch (HMMER V 3.3.1) to query the 120 HMMs against genes present in all ecologies (1,864 and 289 genes from the 30% and 50% gene catalogs respectively).Then we counted the number of HMMs that significantly aligned to the consensus genes (E-value < 0.001 and domain E-value < 0.001). Hmmsearch parameters are --noali, --tblout <OUT_FILE>, --domtblout <OUT_FILE>, -E 0.001, --domE 0.001.

### Software Availability

The figure generation code is located in our GitHub repository, located at https://github.com/kosticlab/pan_ecological_gene_universe.

## Data Access

Data generated in this study is hosted by Figshare and accessible at http://www.microbial-genes.bio. This resource additionally includes files to ensure the reproducibility of our results, specifically the data needed to run the figure generation code in our GitHub repository (see *Software Availability*). A STORMS (Strengthening The Organizing and Reporting of Microbiome Studies) checklist (40) is available at DOI: 10.6084/m9.figshare.20293746

## Acknowledgements

This work was supported in part by Oracle Cloud credits and related resources provided by the Oracle for Research program. This research was additionally supported by the National Institutes of Health NIDDK (T32 DK110919), NIAID(R01AI127250), the American Diabetes Association (ADA) Pathway to Stop Diabetes Initiator Award #1-17-INI-13, and a Smith Family Foundation Award for Excellence in Biomedical Research. We thank Harvard Research Computing for providing compute resources for this work.

## Author contributions

B.T.T. and S.Z. conceived the project. B.T.T. and S.Z. wrote the project code pipeline, conducted the analysis, generated the figures, and wrote the initial draft of the paper. A.D.K. and C.J.P. advised on project progress, statistical methodology, microbiome analyses, and assisted in editing the paper.

## Supplementary Figures and Tables

**Fig. S1: Permutation test of data in** Figure 1. Randomizing the gene-ecology mapping, we recomputed the subsampling analysis in Figure 1 comparing the gene content of different ecologies. A) Hierarchical clustering on the gene overlap between ecologies as a function of iterative sampling. Each cell represents the average number of genes shared between two ecologies (on rows and columns) after 50 random samplings. Cell color is in units of number of genes. Color of text corresponds to broader ecology class. B) The average number of genes unique to (found only in) a given ecology. This number corresponds to the values in the diagonal.

**Fig. S2: Results of our unsupervised, gene-level clustering analysis.** A) Output of our UMAP analysis displaying all seven clusters we identified. Cluster composition corresponds to color. B-C) Using the same color scheme as in A) for the bars, the proportion of samples in different ecologies and with different disease states in our identified clusters. D) Top 10 enriched genera, colored by phyla, in each cluster. E) Distribution of westernized vs non-westernized samples across clusters. F) Distribution of age categories across clusters.

**Fig. S3: Robustness of unsupervised clustering.** We show a series of parameters (described on rows and columns) used to generate UMAP output, as well as the number of clusters generated, silhouette score, and number of unclustered samples. The number of neighbors (10, 15, 20, and 30) and the minimum distances (0.1, 0.25, and 0.5) used to construct the UMAPs are provided on the columns and rows. We also varied the minimum size of clusters. UMAPs with minimum cluster size of 50, 75, and 100 can be seen in a, b, and c respectively. The red box illustrates the chosen UMAP seen in Fig. S2.

**Fig. S4: Potential biases due to depth of sequencing in cluster analysis.** Plot depicts the average number of genes across all samples in all seven clusters identified by unsupervised clustering.

**Fig. S5: Taxonomic enrichment in clusters.** Based on taxonomic annotations of genes, the top 10 enriched phyla, classes, and genera in each of the seven clusters depicted in Figure 2.

**Fig. S6: Functional enrichment in clusters.** The top 25 enriched protein products in each of the seven clusters depicted in Figure 2.

**Fig. S7: Results of sub-clustering analysis.** Rows correspond to the named cluster (with the gray inset showing the original shape of the cluster in Figure 2). Columns correspond to the color scheme of points by ecology, age, and westernization.

**Fig. S8: Sub-clustering of human microbiome clusters colored by health status.** The sub-clustering carried out in Fig. S7 on the four clusters containing human gut microbiome data, but colored by if an individual were healthy or diseased.

**Fig. S9: Number of phyla annotated to proteins enriched in ecological groups.** Companion figure to 3C, 3D, and 3E. Depicts the number of phyla annotated to the top 25 proteins with significantly greater abundance in a) environment ecologies b) gut and environmental ecologies, and c) human gut and non-human gut ecologies.

**Fig. S10: Genes abundant in human body sites.** Figure shows genes that were abundant (in our 422 external samples) in at least 1 human body site compared to non-human ecologies that were not from gut microbiome samples and any overlap therein.

**Fig. S11: Conserved gene analysis, 50% sequence identity.** Companion figure to 6B. Depicts the intersections between the 289 ecologically conserved genes identified at 50% sequence identity and the bac120 GTDB genes.

**Table S1: Sample metadata and summary statistics.** File containing database summary information (e.g. N50s, sample IDs, study IDs, number of genes per sample) as well as sample metadata (e.g. disease or westernization status).

**Table S2: Number of unique functional annotations.** File containing the numbers of unique protein annotations, COG annotations, and EC numbers in the 117,629,181 non-redundant genes.

**Table S3: Metadata for 422 additional samples.** The metadata for the samples used in our abundance calculations of genes identified as being prevalent in different ecological groupings.

**Table S4: Pan-ecologically conserved gene metadata.** File containing the functional annotations and mapping to GTDB genes for the 1,864 pan-ecologically conserved gene sequences identified in our analysis.

